# Mapping and Engineering Antibody-Antigen Interaction Landscapes for Systematic Affinity Enhancement

**DOI:** 10.1101/2024.05.25.595884

**Authors:** Changju Chun, Byeong-Kwon Sohn, Ji Hye Jo, Jiyu Lee, Booyoung Yu, Minkyung Baek, Tae-Young Yoon

**Affiliations:** School of Biological Sciences, Seoul National University, Seoul 08826, South Korea; Institute of Molecular Biology and Genetics, Seoul National University, Seoul 08826, South Korea; Department of Biomarker Discovery, PROTEINA Co., Ltd, Seoul 08826, South Korea

**Author notes:** These authors equally contributed to this work (Changju Chun, Byeong-Kwon Sohn, Ji Hye Jo).

## Abstract

Antibodies, crucial in adaptive immunity, recognize antigens through specific interactions facilitated by Complementarity Determining Regions (CDRs), diversified via Variable-Diversity-Joining (VDJ) recombination. Traditional antibody development, limited by the scope of animal models and phage display libraries, captures a fraction of the potential antibody-antigen interactions. This underscores a gap in understanding antibody specificity and the relationship between antibody sequence and binding affinity. Here we introduce an approach using the Single-Protein Interaction Detection (SPID) platform, repurposed to systematically map local landscapes of antibody-antigen interactions with unprecedented depth and speed, aiming to rival the precision of methods like Surface Plasmon Resonance (SPR) and Bio-Layer Interferometry (BLI) while significantly boosting throughput. By editing CDR sequences and measuring effects on dissociation constants, we elucidated pathways for optimizing antibody affinity, enhancing predictive models for interactions. Our findings demonstrate the capability of the SPID platform to characterize thousands of variants weekly, offering a deeper insight into antibody-antigen interactions and advancing antibody development with finely-tuned affinities.

---

Antibodies are crucial components of adaptive immunity, selectively recognizing and binding to foreign proteins through specific protein-protein interactions that form antigen-antibody complexes [1–3]. Antibodies achieve sequence diversity primarily through Variable-Diversity-Joining (VDJ) recombination, which constructs the Complementarity Determining Regions (CDRs), enabling antibodies to bind a vast array of antigens with varying affinities [4]. Each antibody features six CDRs, each composed of roughly ten polypeptides, distributed across its heavy and light chains [5, 6].

Historically, antibody development has depended on the immunological responses of animal models or on diverse antibody libraries, such as those displayed on bacteriophages [7–9]. These traditional methods essentially probe the antibody-antigen interaction space. The diversity of antibody sequences, derived from B cell sequencing, exceeds 10^12^ [10], and the scale of bacteriophage screenings can surpass 10^11^ [11]. Nevertheless, these figures are starkly modest compared to the potential combinations of CDR sequences, estimated at approximately 20^60^. This disparity indicates that the conventional methods result in a very coarse sampling of the antibody-antigen space, underscoring a fundamental gap in our understanding of the complex relationship between antibody sequence and antigen-binding affinity. In addition, traditional methods often yield only the end products of selection [9], obscuring the evolutionary pathways of selection and further reducing the ability to systematically explore the antibody-antigen binding landscape.

There is an escalating demand for systematic sampling of the antibody-antigen space, surpassing the capabilities offered by the traditional methods. By introducing extensive variations to the CDR sequences and analyzing the resultant effects, we can begin to delineate how local interaction landscapes shape antibody-antigen interactions. Mapping these landscapes for numerous antibodies can lead to illumination of the broader contours of the antibody-antigen interaction space. This would offer invaluable opportunities for novel deep learnings that are aimed at deciphering the underlying patterns governing the antibody-antigen interactions. This approach promises to enable more rational and efficient optimization of lead antibody candidates.

In this study, we have developed an experimental streamline utilizing the Single-Protein Interaction Detection (SPID) platform, previously established for sensitive detection of protein-protein interaction (PPI) complexes in clinical specimens [12–14]. This adaptation of the SPID platform enables us to more deeply sample local landscapes of specific antibody-antigen interactions. Our objective was to achieve the same level of precision provided by current gold standards such as Surface Plasmon Resonance (SPR) and Bio-Layer Interferometry (BLI), enabling us to determine precise dissociation constants (*K*_D_) for individual antibody-antigen interactions, rather than relying on arbitrary and relative affinity scores [15, 16]. Concurrently, we explored whether our method could handle the high throughput analysis of over 3,000 different antibodies (with varied CDRs) per week. This capability positions our approach uniquely, allowing for rapid navigation through local landscapes of antibody-antigen interactions at a scale of approximately 10,000 variants per week (when employing three pipelines in parallel). The data resolution is maintained at the level of dissociation constants, which can be directly translated into thermodynamic quantities such as free energy differences. Therefore, the resulting data sets from our method can be seamlessly integrated without the need for complex recalibration, forming comprehensive data sets that reveal the extensive landscape of antibody-antigen interactions. To demonstrate the utility of these local landscapes, we mapped how interactions between HER2 and Trastuzumab altered upon varying each residue in the CDRs of Trastuzumab to all 20 possible amino acids [17]. The resultant map highlighted clustered regions of affinity-enhancing single-residue variants. By combining these variants, we successfully engineered Trastuzumab variants with significantly enhanced affinities in the sub-nanomolar range.

## Results

### Determination of *K*_D_ values for antibody-antigen interactions with single-molecule resolution

To generate large-scale data for antibody-antigen interactions, we employed single-chain Fab (scFab) proteins, known to effectively replicate the full complexity of antibody-antigen interactions (Fig. 1a). Due to their smaller size, scFab proteins typically exhibit higher expression levels than full IgGs . We selected the interaction between trastuzumab and HER2 as a model system [13, 17]. We engineered the heavy and light chains of trastuzumab into an scFab vector with a polypeptide linker of 60 amino acids long, tagging the C-terminal end with a red fluorescent protein (RFP, specifically mScarlet). Successful expression and secretion of these scFab proteins into the culture media were confirmed, where secreted RFPs fully linked to scFab proteins with no significant cleavage between the heavy and light chains or between scFab and the RFP tag (Fig. 1b and Supplementary Fig. 1). We directly immobilized these RFP-labeled scFab proteins onto the surface of a SPID imaging chip using anti-RFP antibodies as there was no mammalian homolog of red fluorescent proteins in both cell culture media and cell-secreted proteins (Fig. 1c). Of note, this direct immobilization was made possible by the uniform poly-ethylene glycol (PEG) layer coated on the SPID imaging chip, which nearly eliminated non-specific adsorption of proteins onto the surface (confirmed with mass-spectrometry imaging; T.-Y.Y. unpublished results), thereby bypassing the need for a purification process. The pull-down number increased linearly with the concentration of scFab in the media. Using antibodies targeting the Fc domain of human antibodies, the pull-down number dropped to background levels, confirming the specificity of the surface pull-down process (Fig. 1d).

**Fig. 1.**
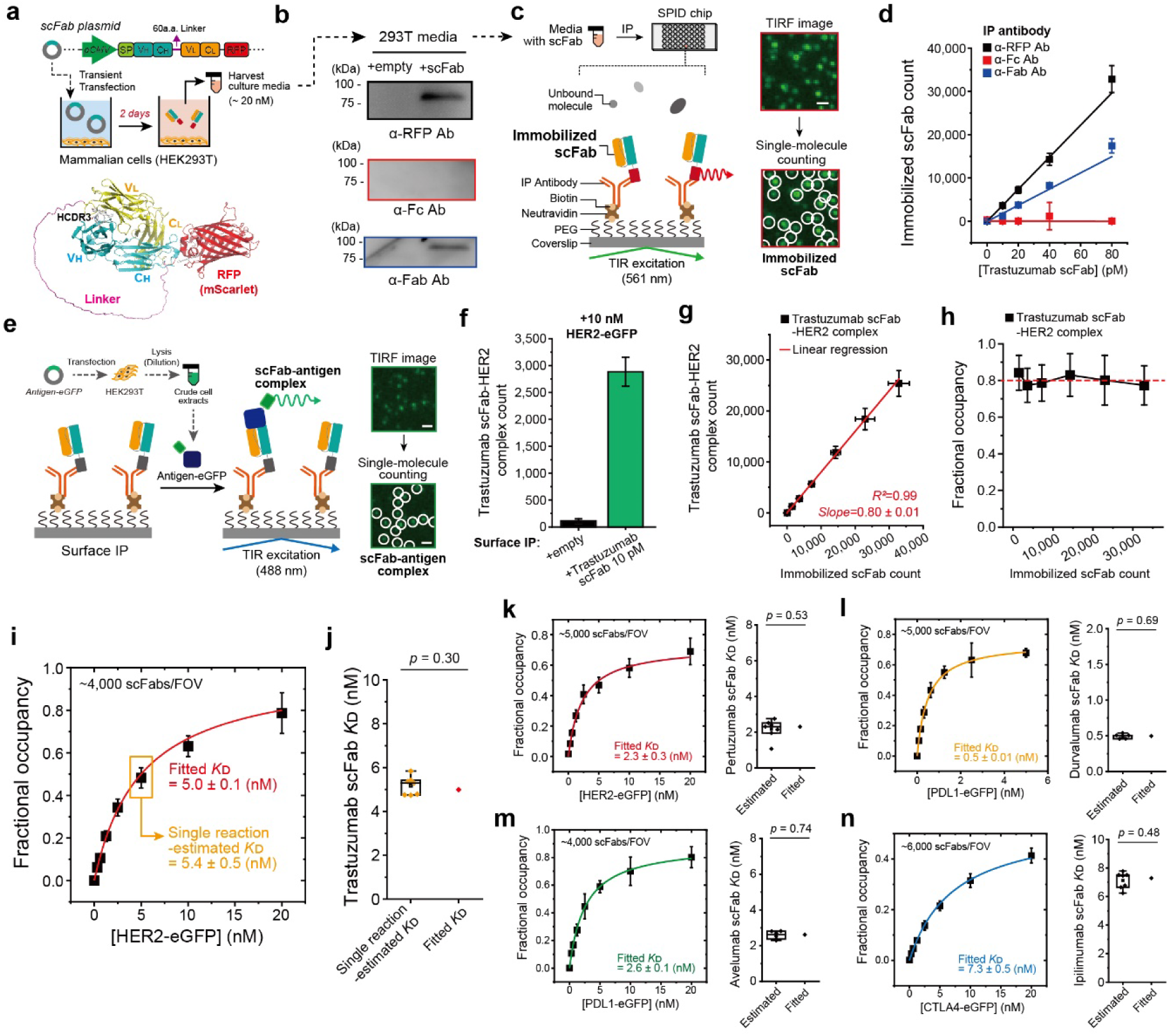
SPID platform allows for quantifying binding affinities between scFabs and antigens. **a**, Schematic for the construct design and expression of Trastuzumab based scFab. **b,** Detection of the secreted Trastuzumab scFabs in cell culture media with Western blotting. **c,** Schematic for the surface immobilization of scFab proteins with SPID platform. scFab proteins in cell culture media were directly immobilized onto the surface of the SPID imaging chip. **d,** Single-molecule count of surface-immobilized Trastuzumab scFabs with indicated IP antibodies. **e,** Schematic for the detection of scFab-antigen interactions with SPID platform. The eGFP-labeled antigen proteins were expressed in HEK293T cells and directly introduced on the surface-immobilized scFabs to form scFab-antigen complexes. **f,** Single-molecule count of Trastuzumab scFab-HER2 complexes. The HER2-eGFP antigens were loaded onto the surface-immobilized Trastuzumab scFabs with anti-RFP antibodies. **g,** Linear increase in the Trastuzumab scFab-HER2 complex count with increasing amounts of surface-immobilized Trastuzumab scFabs. **h,** Remaining of fractional occupancies with increasing amounts of surface-immobilized Trastuzumab scFabs. **i,** Binding curves for HER2-eGFP to Trastuzumab scFab used for *K*_D_ fitting. **j,** Comparison of K_D_ values for the Trastuzumab scFab-HER2 interaction obtained by single reaction-estimated and fitted manners (Two-sided two-sample t-test). **k-n,** Development of various scFab-antigen pairs for SPID platform. Binding curves for eGFP-labeled antigen proteins to its pair scFabs used for *K*_D_ fitting. The *K*_D_ values obtained by single reaction estimated and fitted manners were compared (Two-sided two-sample t-test). (k) PertuFab-HER2, (l) DurvalFab-PDL1, (m) AvelFab-PDL1, (n) IpiliFab-CTLA4.

For full-length human HER2 antigens, we fused an enhanced green fluorescent protein (eGFP) at the C-terminal end and expressed these eGFP-labeled antigens in HEK293T cells (Fig. 1e). Our prior experiments showed that lysis with specific non-ionic detergents effectively preserves the conformation of this integral membrane protein with a single transmembrane domain [13]. After lysis, the crude extracts were diluted to achieve a final HER2-eGFP concentration of 10 nM and added to the reaction chamber, facilitating reconstitution of the antibody-antigen binding interaction on the surface of the SPID imaging chip (Fig. 1e,f). Stable formation of antigen-antibody complexes was visualized as diffraction-limited spots using a total internal reflection (TIR) fluorescence microscope under excitation at 488 nm. Without immobilization of scFabs (using an anti-RFP antibody), the count of scFab-antigen complexes diminished to baseline levels, further supporting the specificity of the antibody-antigen reaction, even though the reaction was reconstituted in lysate environments (Fig. 1f).

Remarkably, the number of antibody-antigen complexes linearly increased up to a point where approximately 35,000 scFab proteins were immobilized per field of view (86×86 µm² provided by the SPID imaging machine), without any signs of saturation (Fig. 1g). This indicated that the fractional occupancy value was consistently maintained when the antibody immobilization number varied by more than 20-fold (Fig. 1g,h). We then varied the antigen concentration while maintaining the antibody immobilization number at approximately 4,000 per field of view (Fig. 1i). The resulting fractional occupancy followed the well-known Hill equation with a Hill coefficient of 1, yielding a dissociation constant of 5.0±0.1 nM, close to the reported values for Trastuzumab based on SPR and cell-based assay (Fig. 1i, red) [18–20]. In addition, we explored whether the dissociation constant could be accurately estimated through a single measurement at one antigen concentration, rather than fitting to the Hill equation across multiple data points (for example, Fig. 1i, yellow box). The distribution of *K*_D_ values estimated from each individual measurement exhibited a narrow range, with a mean value of 5.2±0.4 nM (Fig. 1j). This consistency hints at the potential for rapid determination of *K*_D_ values via single measurements. This efficiency is achieved by precisely quantifying the scFab pull-down number, antigen concentration, and resultant antigen binding number, leveraging the single-molecule resolution afforded by the SPID platform.

To test the general applicability of this method, we cloned four more blockbuster antibody drugs (Pertuzumab, Durvalumab, Avelumab and Ipilimumab) into scFab vectors and reconstituted their interactions with antigens on the SPID imaging chip following the same procedure [21–24]. All resultant antibody-antigen complex counts precisely followed the Hill equation, leading to the ascertainment of respective *K*_D_ values of 2.3±0.3 nM, 0.5±0.01 nM, 2.6±0.1 nM, and 7.3±0.5 nM, closely aligning with reported values for each antibody drug (Fig. 1k-n). We also confirmed that the scFab version of Pertuzumab can bind to single full-length HER2 proteins simultaneously with Trastuzumab, consistent with their disparate binding sites on HER2 (Supplementary Fig. 2). Moreover, when *K*_D_ values were assessed based on each data point, all the results showed narrow distributions, centered around those determined from fitting the entire Hill curve. These data reaffirm the feasibility of *K*_D_ determination based on single reaction measurements (Fig. 1k-n).

### Mapping local landscapes of antibody-antigen interactions with increased speed

Based on the demonstrated capability of the SPID platform to characterize antibody-antigen interactions via single sets of measurements, we sought to develop an experimental procedure to rapidly generate and assess thousands of different antibodies. A critical focus was on the precise editing of CDR sequences in individual antibodies and the direct measurement of absolute *K*_D_ values, rather than merely assigning arbitrary relative affinity scores.

In the first step of our experimental streamline, we introduced sequence variations into the CDR sequences of antibodies. We used the scFab version of Trastuzumab, as shown in Figure 1, and targeted the sequence of its HCDR3. We introduced silent mutations, which did not alter the amino acid sequence but created restriction recognition sites for *XbaI* and *KpnI* at the 5’ and 3’ positions of HCDR3, respectively (Fig. 2a). Concurrently, we removed all other restriction sites from the scFab vector and inserted five stop codons in the middle of HCDR3 to ensure that any undigested vectors remaining after enzyme digestion would not contribute to scFab production (Fig. 2b).

**Fig. 2.**
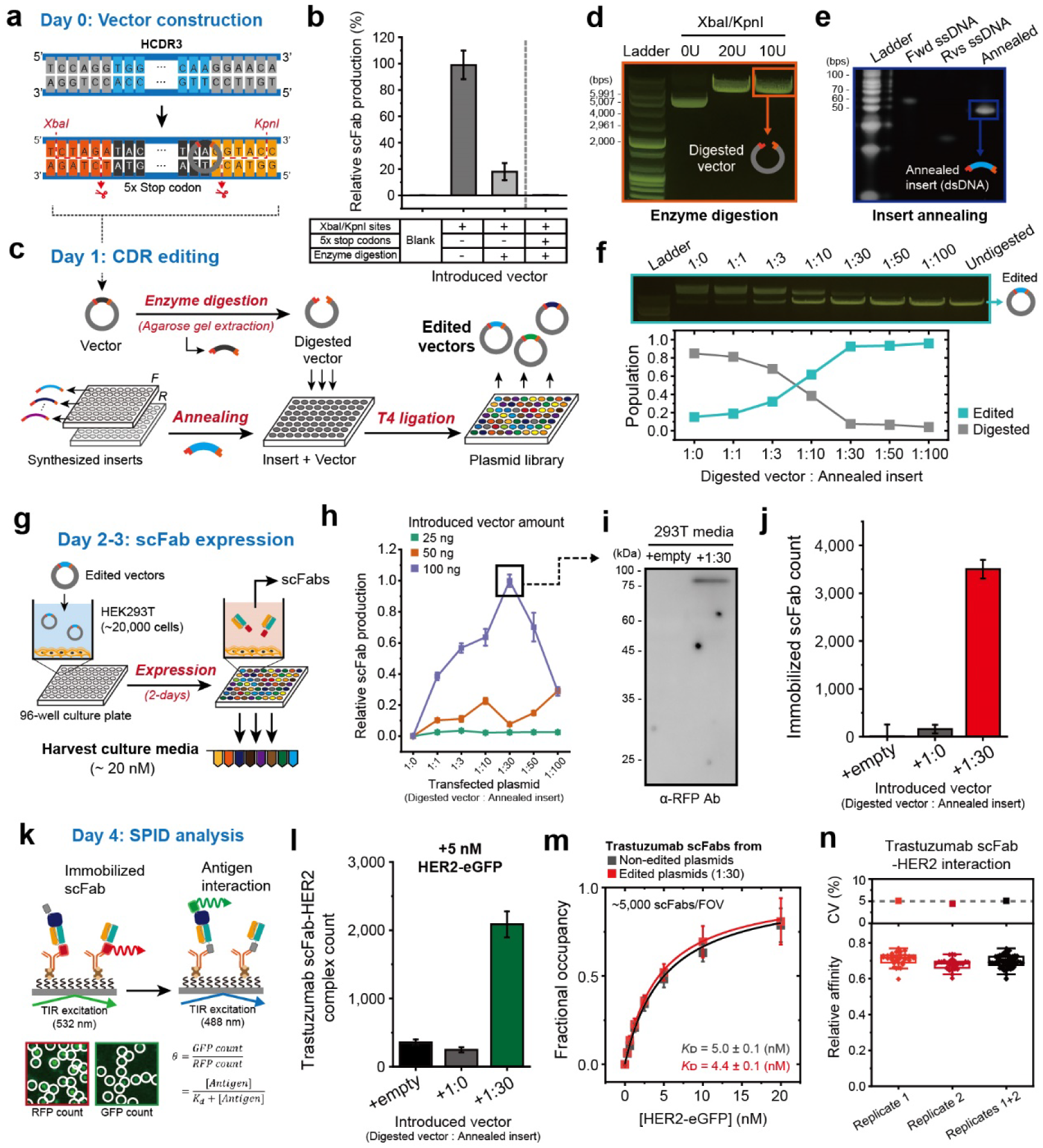
Application of SPID to generate scFabs and determine *K*_D_ values with increased speed. **a**, Construction of the Trastuzumab scFab encoding expression vector for rapid HCDR3 editing. **b,** Complete inhibition of scFab production by introducing of five stop codons in the middle of the HCDR3. **c,** Schematics for the high-throughput *in vitro* CDR editing. Synthesized ssDNA inserts were prepared the day before CDR editing. **d,** Comparison of digestion efficiency using restriction enzymes. Trastuzumab scFab vector construct was digested with *KpnI* and *XbaI* simultaneously. >99% purity of digested vectors were obtained. **e,** Comparison of annealing efficiency between forward and reverse ssDNA insert fragments. >99% purity of annealed DNA insert fragments were obtained. **f,** Comparison of *in vitro* ligation efficiency according to the ratio of insert fragment to digested vector. **g,** Schematics for scFab expression after CDR editing. **h,** Expression levels of Trastuzumab scFabs from 293T cells after 2 days of expression according to the introduced ligated plasmid. **i,** Validation of secreted Trastuzumab scFabs by using Western blotting. **j,** Detection of the secreted Trastuzumab scFabs using SPID. **k,** Schematics for affinity measurement of scFabs using SPID. **l,** Detection of specific interaction between secreted Trastuzumab scFabs and HER2-eGFP proteins using SPID. 10 nM of HER2-eGFP were loaded onto immobilized scFab in panel (**j**). **m,** Comparison of the binding curves of Trastuzumab scFab-HER2 interactions according to the scFab production methods. **n,** Reproducibility for binding affinities of Trastuzumab scFabs. 30 distinguished CDR editing to rescue the original HCDR3 sequence were performed in each trial. Intra-chip CVs were displayed.

We digested the recombinant vector with *XbaI* and *KpnI*, followed by separation through agarose gel extraction (Fig. 2c,d). We selected an insert encoding the original HCDR3 sequence, which did not contain stop codons, to rescue antigen affinity if CDR editing were successful. The inserts, initially synthesized as single-stranded DNAs (ssDNAs) complemented as forward and reverse directions, were annealed by heating and progressive cooling (Fig. 2c,e). After the complementary double-stranded DNA (dsDNA) formation, the annealed inserts were combined with the digested vectors at different molar ratios, and connected with T4 ligase to find the optimal molar ratio (Fig. 2c,f). This procedure typically yielded 400 ng of the final vector with the swapped HCDR3 sequence. This series of processes, while bypassing any steps involving micro-organisms (typically *E. coli*), consists solely of *in vitro* reactions. Thus, by increasing the concentrations of reagents, we were able to minimize stochastic variations in these steps.

We transfected the edited vectors into a culture of HEK293T cells and induced scFab production and secretion for two days (Fig. 2g). By titrating the starting mass of the vector, we discovered that a minimal mass of 100 ng was sufficient for scFab production at levels (up to 5 nM) suitable for characterization on the SPID platform (Fig. 2h,i). The scFab production exhibited a biphasic pattern with a production peak at a vector-to-insert ratio of 1:30, indicating that excess inserts interrupted effective transfection of the edited vectors on to HEK293T cells.

The SPID platform’s compatibility with complex biological extracts allowed for direct utilization of the culture media containing secreted scFabs. As demonstrated in Figure 1, we immobilized the scFab proteins on the surface using anti-RFP antibodies, minimizing non-specific adsorption of various proteins. In this trial, under conditions of 100 ng vector, a 1:30 vector-to-insert ratio, and a 2-day culture, we were able to immobilize approximately 3,500 scFab proteins per field of view from 10 pM of diluted samples (Fig. 2j). We then added the eGFP-tagged HER2 antigen at 5 nM to the SPID imaging chip and observed the formation of approximately 1,750 of net antibody-antigen complexes on average, corresponding to a fractional occupancy of ∼50 % and a *K*_D_ value of 5.1±0.6 nM, very close to the *K*_D_ value determined before CDR editing (Fig. 2k,l). Meanwhile, only the digested vectors, indicated as a vector-to-insert ratio of 1:0, did not produce scFab as well as did not interact with HER2 (Fig. 2j,l). Finally, the *K*_D_ of trastuzumab scFab produced from edited plasmid showed at almost the same level as that produced from non-edited plasmid (Fig. 2m).

We repeated the entire procedure from insert synthesis to scFab production across 30 independent trials, resulting in 30 cell media samples containing scFab (i.e., 30 biological replicates) (Fig. 2n). We subsequently conducted two technical replicate measurements for each antibody-containing medium, totaling 60 measurements. The *K*_D_ values from these measurements showed a very narrow distribution with coefficients of variation of less than 5%, indicating a remarkable level of reproducibility and stability in this experimental streamline leveraging the SPID platform. Moreover, the entire process from CDR editing to *K*_D_ determination could be completed within five business days, suggesting the potential of the streamline described here that can generate more than a thousand data points within a week.

### SPID reveals local landscapes of the Trastuzumab-HER2 interactions

Leveraging the experimental procedures established previously, we embarked on generating a comprehensive data set by systematically substituting each amino acid in three different CDRs (HCDR2, HCDR3, and LCDR2) and a partially structured framework 3 (F3) directly linked to HCDR2 of Trastuzumab with all 20 naturally occurring amino acids (Fig. 3a). This endeavor resulted in the examination of 520 variants, given that there were 26 residues in the targeted domains. In the resulting heatmap, where stronger and weaker affinities were denoted by red and blue, respectively, and the original *K*_D_ value by white, we observed a predominance of white with interspersed red and blue points, signifying mostly subtle changes in affinity (Fig. 3a). Importantly, this expansive data set demonstrated that many sequence variations were neutral, having minimal impact on binding affinity (Fig. 3b).

**Fig. 3.**
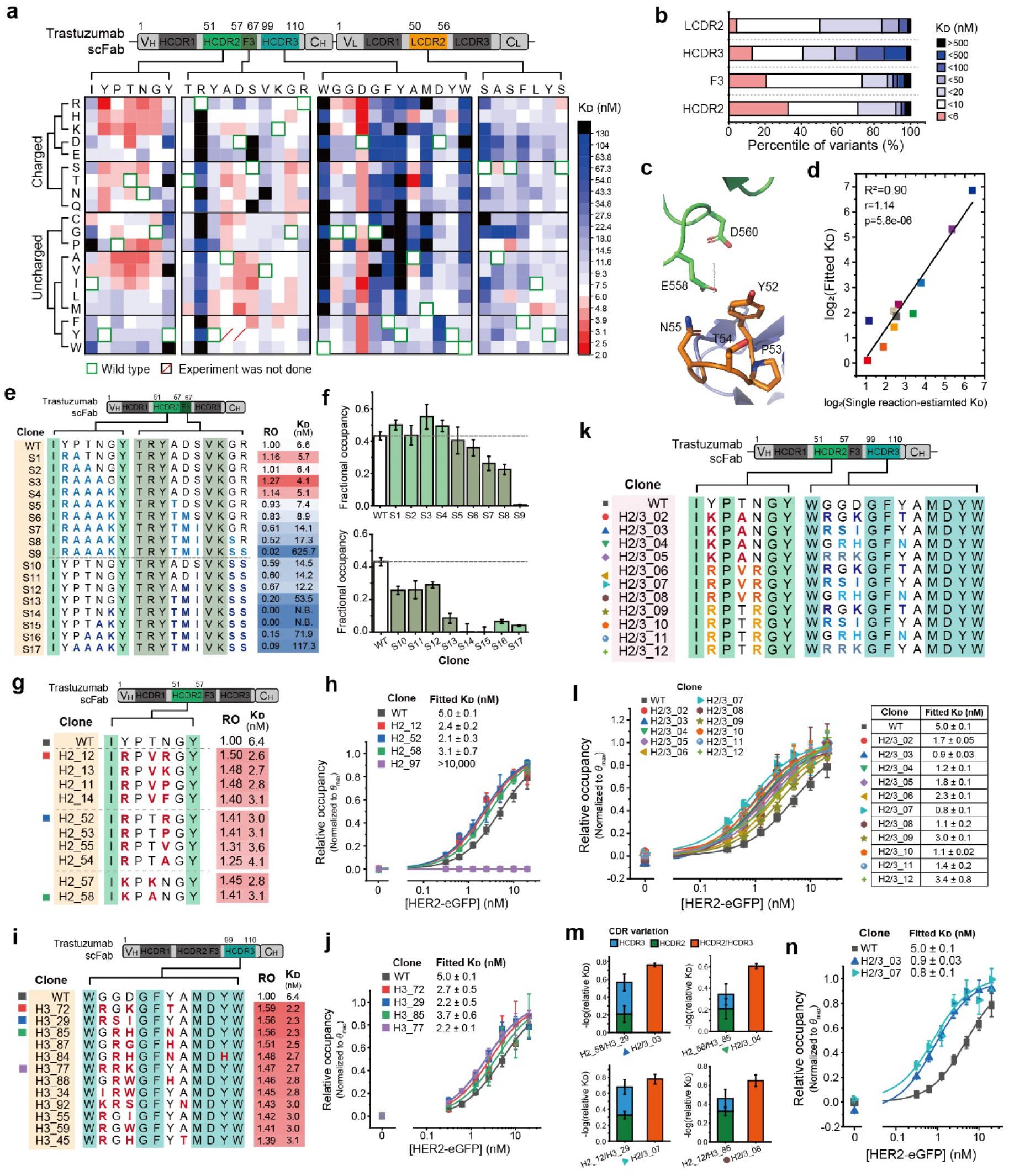
SPID reveals a local landscape of antibody-antigen interactions. **a,** Screening the *K*_D_ changes of Trastuzumab scFab-HER2 interactions from single-residue variations in CDRs (HCDR2, HCDR3, and LCDR2) and a framework 3 (F3). **b,** Distribution of *K*_D_ changes for Trastuzumab scFab. **c,** Structure of Trastuzumab Fab-HER2 complex (PDB id: 1N8Z). HER2 and Trastuzumab HCDR2 were depicted as green and orange, respectively. **d,** Correlation between the single reaction-estimated *K*_D_s by SMASH and fitted *K*_D_s from 11 different Trastuzumab scFab variants obtained by conventional cloning method. **e,** Progressive combination of single-residue variations with lowest *K*_D_ values in HCDR2/F3 to increase the length. **f,** Comparison of the fractional occupancies in panel (**e**). **g,** Changes of *K*_D_s in Trastuzumab scFab with multiple-residue variations in HCDR2. **h,** Binding curves for Trastuzumab scFab variants and HER2. **i,** Changes of *K*_D_s in Trastuzumab scFab variants with multiple-residue variations in HCDR3. **j,** Binding curves for Trastuzumab scFab variants and HER2. **k,** Combination of HCDR2 and HCDR3 variants with high-binding affinities. **l,** Changes of *K*_D_s in Trastuzumab scFab with multiple-residue variations both in HCDR2 and HCDR3. **m,** Synergistic analysis for cross-combination of HCDR2 and HCDR3 variants. **n,** Determination of Trastuzumab scFab variants with enhanced binding affinities for HER2. (**e,g,i,k**) The altered residues are indicated. RO: relative occupancy. (**h,j,l**) The Trastuzumab scFab variants were cloned with conventional methods.

The heatmap vividly illustrated the nuanced landscape of the Trastuzumab-HER2 interaction. While most amino acid substitutions led to minor affinity alterations, several changes in the HCDR3 region significantly reduced or nearly abolished binding, highlighted by numerous blue and black points (Fig. 3a,b). This pattern underscores the critical role of HCDR3 in the Trastuzumab-HER2 interaction and suggests a high degree of evolutionary optimization, where even minor modifications could drastically decrease affinity. Notably, virtually all variations at position D102 led to enhancement of binding affinity, standing out as an anomaly.

In a sharp contrast, the HCDR2 region displayed many clusters of red points, indicating potential for further affinity optimization. Analysis of the variations relative to *K*_D_ values showed that substitutions in HCDR2 frequently yielded lower *K*_D_ values compared to the original sequence (Fig. 3a,b). This trend was particularly pronounced when residues between positions 52 and 56 were substituted with positively charged residues (R, H, K) (Fig. 3a), which likely enhanced affinity due to increased electrostatic interactions with the negatively charged residues E558 and D560 located in loop 1 domain of HER2 (Fig. 3c). On the contrary, modifications of many of these residues to either polar or hydrophobic types also improved affinity, underscoring the complexity of the Trastuzumab-HER2 interaction and suggesting that multiple routes may exist to augment affinity towards the HER2 antigen. Additionally, replacing D62 and S63 in F3 with hydrophobic residues significantly increased affinity, pointing to enhanced hydrophobic interactions at the binding interface (Fig. 3a). Notably, the substitutions of small residues such as G56, A61, and G66 across HCDR2 and F3 to virtually any other residue type generally maintained or increased affinity, highlighting their versatile role in fine-tuning the interaction.

To validate the accuracy of our *K*_D_ estimations based on single measurements, we selected 11 single-residue variants spanning different affinity ranges for further analysis (Fig. 3d). These antibodies were produced in substantial quantities via traditional cloning methods, and their *K*_D_ values were reassessed by titrating antigen concentrations and fitting data to the complete Hill equation as done in Figure 1i. The resulting *K*_D_ values from this detailed analysis correlated strongly with those from our initial screenings in Figure 3a (R²=0.90), reaffirming the reliability of our high-throughput approach in determining dissociation constants through single measurement protocols.

### Systematic engineering toward high-affinity antibodies based on PPI landscapes

Informed by the local landscape of the Trastuzumab-HER2 interaction, we sought to systematically combine multiple variations to engineer Trastuzumab variants with enhanced affinity for the HER2 antigen. Given the vast size and complexity of the entire antibody-antigen interaction space, our strategy focused on integrating single-residue variants that demonstrated affinity enhancement (red points in the heatmap), avoiding arbitrary combinations. Specifically, we identified 117 variants in HCDR2/F3 and 53 in HCDR3 of Trastuzumab that enhanced affinity in our single variant screening (Supplementary Fig. 4).

Our efforts concentrated on combining affinity-enhancing variations in HCDR2/F3. We maintained Y57, R59, and V65 as constants because alterations at these positions often completely nullified Trastuzumab’s binding affinity (Fig. 3a). We identified variations between positions 52 and 67 that resulted in the lowest *K*_D_ values from the 20 variants examined per residue, and progressively combined these variations to increase their length (Fig. 3e). Apart from the initial variation Y52R, which exhibited a *K*_D_ of 2.4 nM, extending this combination in an indiscriminate manner only triggered antagonistic interactions, consistently reducing the overall affinity and suggesting a deviation from local maxima in the antibody-antigen interaction landscape (Fig. 3e,f).

Instead, our empirical approach revealed that strategic inclusion of one residue from the original Trastuzumab sequence as spacers between the affinity-increasing variations optimized the enhancements (Supplementary Fig. 4). We particularly concentrated on positions 52 to 56 in HCDR2, where alterations to either positively charged, polar, or hydrophobic residues consistently yielded clusters of red points on the heatmap (Fig. 3g). Combinations of residues with disparate electrical properties, such as IRPVRG, IRPTRG, and IKPANG, emerged from our exploratory methodology, which reduced *K*_D_ to 2.6, 3.0, 3.1 nM, respectively (Fig. 3g). The enhanced affinities observed with these combinations were rigorously confirmed through comprehensive measurements across multiple antigen concentrations and precise fitting to the full Hill equation (Fig. 3h).

Similarly, we applied a strategy in combining single affinity-enhancing variants for HCDR3 (Supplementary Fig. 4). We established residues 99W, 103G, 104F, and 107M-110W as fixed points, as alterations here significantly reduced binding (Fig. 3i). We then targeted the segments 100G to 102D and 105Y to 106A for variations, incorporating strategic spacers to reflect the original heatmap (Fig. 3a). This method yielded several combinations that consistently enhanced affinity, confirming the effectiveness of our approach in utilizing the PPI landscape to guide the development of antibodies with significantly improved binding characteristics (Fig. 3i,j).

Finally, we sought to determine if the advantageous combinations identified for HCDR2 and HCDR3 could be merged to further enhance the affinity of Trastuzumab. We selected three HCDR2 variant clones—H2_58, H2_12, and H2_52—each demonstrating the lowest *K*_D_ values among those tested in each cluster, alongside four HCDR3 variants—H3_72, H3_29, H3_85, and H3_77—that also exhibited the lowest *K*_D_ values in their group (Fig. 3g,i). Given the presence of more than 30 residues separating the variations in HCDR2 and HCDR3, which act as natural spacers, we directly combined these clones without adding additional buffer spacers (Fig. 3k,l). Remarkably, 4 out of the 11 possible cross-combinations between HCDR2 and HCDR3 variant clones (one combination was not examined due to experimental constraints) demonstrated synergistic interactions, resulting in *K*_D_ values lower than those predicted by simple additive effects (Fig. 3l,m). However, four of the cross-combinations yielded *K*_D_ values higher than expected (Supplementary Fig. 5). Notably, the clone H2/3_03 and H2/3_07, which are the combination of clones H2_58 or H2_12 with H3_29, achieved sub-nanomolar *K*_D_ values, indicating significant sixfold enhancements in affinity compared to the original Trastuzumab (Fig. 3n).

## Discussion

Antibody optimization traditionally relies on extensive libraries displayed on bacteriophage, *E.coli*, or yeast membranes. These methods typically yield only the end products of selection, with detailed selection routes often remaining obscure. Partial reconstruction of the selection history is possible through techniques like FACS and/or next-generation sequencing, yet precise assignment of *K*_D_ values often remains challenging, resulting in datasets with limited depth and integrability. In contrast, methods determining antibody-antigen structures provide atomic-level interaction details but suffer from low throughput, even with advancements in cryo-electron microscopy. Consequently, SPR and BLI have remained the methodologies of choice for rigorous characterization of antibody-antigen interactions. Despite the availability of high-throughput versions commercially, these techniques still require relatively large quantities of antibodies (over 1 ng) and multiple measurements at different antigen concentrations. Our own attempts to characterize 300 antibodies within a month, using 1 ng of each antibody, encountered failure rates exceeding 50% (Supplementary Fig. 3).

Our principal aim was to determine whether higher throughput could be achieved without compromising the accuracy level provided by SPR and BLI. The SPID platform characterizes antibody-antigen interactions under equilibrium conditions, and leverages its single-molecule resolution to precisely quantify the numbers of immobilized antibodies and bound antigens per field of view. Given the antigen concentration used, the *K*_D_ values can be directly assessed. Notably, we found that single-reaction measurements suffice for accurate *K*_D_ estimations, comparable to those obtained through comprehensive fitting to the Hill equation, albeit with marginally increased error tolerances.

As the SPID platform requires only 10 pg of antibodies for characterization, we devised an experimental pipeline to produce antibodies with variant CDRs in the small quantities needed, and to subsequently characterize their differentiated interactions with target antigens. We developed a straightforward CDR editing process using only *in vitro* reactions with restriction enzymes and T4 ligases. By increasing enzyme concentrations and adjusting the molar ratios of backbone vectors to DNA inserts, we circumvented the need for sequencing to confirm CDR edits. While the process described here allowed for editing of only one CDR at a time, we have since innovated a method to simultaneously edit multiple CDRs in a single-pot reaction, avoiding gel extraction steps (B.-K.S. and T.-Y.Y., unpublished data). Remarkably, 60 repetitive trials produced virtually identical *K*_D_ values, confirming the accuracy of the CDR editing and the robustness of the SPID platform in facilitating consistent, large-scale data generation. Collectively, these streamlined procedures enabled us to determine *K*_D_ values for over 3,000 different antibodies within a single week.

With our streamlined protocol, we mapped the local interaction landscape between Trastuzumab and HER2 by substituting every residue in HCDR2, HCDR3, LCDR2, and F3 with all 20 natural amino acids. We identified several residues as critical to the interaction, where variations significantly altered the *K*_D_ values. In contrast, changes in other residues resulted in less than a threefold modification of the *K*_D_ values, indicating that these residues contribute incrementally and collectively to the Trastuzumab-HER2 interaction. This suggests potential for fine-tuning the antibody’s affinity through targeted residue editing. However, the distribution of critical versus ancillary residues was not uniform across the CDRs, underscoring the differentiated roles of each CDR in mediating binding. Particularly, many residues in HCDR3 abolished HER2 binding when varied, confirming their pivotal role in the interaction. Conversely, several single-residue variations in HCDR2 enhanced affinity, highlighting substantial opportunities for optimizing the HCDR2 sequence.

In our quest for systematic enhancement of Trastuzumab’s affinity, we explored combining single residue variations that individually increased HER2 affinity. Surprisingly, consecutive, indiscriminate combinations of these variants frequently led to antagonistic interactions, significantly reducing affinity. We discovered that interspersing one or two residues from the original Trastuzumab sequence between these affinity-enhancing variants provided a more effective strategy for engineering CDR sequences with increased affinities. The rationale for why certain combinations outperform others remains elusive, even with detailed knowledge of the Trastuzumab-HER2 binding structure. Moreover, the final combinations of HCDR2 and HCDR3 variant clones resulted in both synergistic and antagonistic interactions. It is noteworthy that the experimental streamline described in this work, including the synthesis of DNA inserts, can be completed within two weeks. This rapid turnaround capability of the SPID platform opens a new pathway for exploring the intricate dynamics of the antibody-antigen interaction space.

## Materials and Methods

### Protein constructions

All cDNAs of variable domains including the heavy and light chains of recombinant scFabs (Trastuzumab, Pertuzumab, Avelumab, Durvalumab, Ipilimumab) were synthesized by Macrogen (South Korea). All cDNAs were cloned into *pCMV* vectors which removed the 18 different restriction enzyme sites to generate scFab constructs. The mScarlet sequences were cloned into the scFab constructs to generate mScarlet-labeled antigen constructs.

The antigen proteins (HER2, PDL1, and CTLA4) were isolated from their respective human cDNA. All cDNAs were cloned into *pCMV* vectors with eGFP sequences to generate eGFP-labeled antigen constructs.

### Cell culture and harvest

HEK293T cells were purchased from American Type Culture Collection (ATCC). HEK293T cells were cultured in DMEM supplemented with 10% (v/v) FBS and 100 μg/ml penicillin/streptomycin. The scFabs and the eGFP-labeled antigen proteins were expressed in the HEK293T cells through transient transfection. The DNA plasmids were introduced into HEK293T cells in a 90mm^2^ cell culture plate. For scFab expression, the cell culture media containing scFabs were collected 2 days after transfection, and cell debris were eliminated by centrifugation. The supernatants were stored at -80 °C by snap freezing. For antigen protein expression, the cells were collected with the scraper in cold DPBS after 24 hours expression. The cell suspensions were centrifuged at 4 °C, and the supernatants were discarded. The cell pellets were stored at -80 °C by snap freezing. All collected cell pellets were resuspended with the lysis buffer including non-ionic detergent and incubated for 30 minutes at 4 °C. After lysis, the cell suspensions were isolated after centrifugation. The fluorescence protein concentrations were quantified with a microplate reader. The supernatants were aliquoted and stored at -80 °C followed by snap-freezing with liquid nitrogen.

### Antibodies

Anti-RFP with biotin conjugation, anti-human IgG Fcγ with biotin conjugation, and anti-human IgG (Fab’)2 with biotin conjugation were used to immobilize the scFabs for SPID as well as used to detect the scFabs by western blotting.

### Western blotting

The cell culture media containing scFabs were heated at 95 °C with SDS containing sample buffer and resolved with 12% polyacrylamide gels. After SDS-PAGE, the PVDF membranes were blocked with 1% BSA in TBST buffer, and the membranes were immunoblotted with respective antibodies described on Antibodies subsection. The protein bands were labeled with streptavidin conjugated with HRP and detected by using an ECL imaging system.

### CDR editing

The plasmids encoding Trastuzumab scFab were digested by reacting two pre-designated restriction enzymes at 37 °C for 2 hours. After enzyme digestion, the mixture selectively obtains only the digested vector from which the CDR encoded region has been removed through an 0.8% agarose-gel electrophoresis and agarose-gel purification system. The forward ssDNA inserts fragment and reverse ssDNA insert fragment encoding the CDR are combined into annealed dsDNA fragment through stepwise cooling. ssDNA and dsDNA fragments were resolved with 18% polyacrylamide gel and detected after SYBR Gold staining. The digested vector and annealed dsDNA fragment are mixed at a ratio of 1:0 to 1:100, and then ligated plasmid is obtained through ligase reaction. The ligated plasmids were introduced into ∼2.0x10^4^ HEK293T cells in a 96-well cell culture plate through transient transfection. After 2 days expression, the cell culture media containing scFabs were collected after centrifugation to eliminate cell debris.

### SPID

SPID was performed using the Pi-Chip and Pi-View system (Proteina, South Korea). 5 μg/ml of NeutrAvidin was loaded into each reaction chamber of the Pi-Chip. After washing the imaging chip with T buffer (Proteina, South Korea), anti-RFP antibodies with biotin conjugation were loaded to each reaction chamber with 1:1,000. Then, cell culture media containing respective scFabs were loaded into each reaction chamber to immobilize the scFabs on the chip surface with 1:500 (typically, 10 pM) and the chip was washed with T buffer. The cell extracts containing eGFP-labeled antigens were then loaded into each reaction chamber. After incubated for 1 hour to reach the binding equilibrium, surface-immobilized scFab-antigen complexes were fixed with F buffer (Proteina, South Korea) and the chip was moved on to Pi-View (Proteina, South Korea) to acquire fluorescence images. Single-molecule fluorescence imaging was performed with two different excitation lasers for 488 nm and 561 nm wavelengths. The emitted photons were collected by sCMOS camera in Pi-View and converted to the single-molecule fluorescence images. From the obtained fluorescence images, the single-molecule spots were determined by Pi-Analyzer software (Proteina, South Korea). The number of surface-immobilized scFabs (RFP signals) and the number of scFab-antigen complex (GFP signals) were converted to fractional occupancy, and then the dissociation constants (*K*_D_s) were calculated by the following equation.

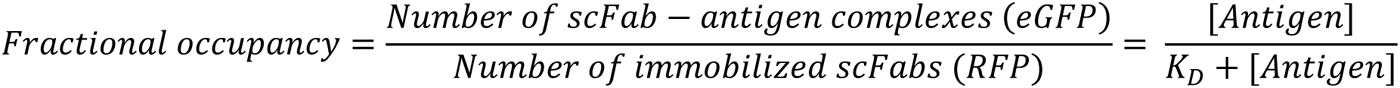

## Author contributions

T.-Y.Y. conceived of and supervised the project. C.C. designed the experiments. C.C, B.-K.S., J.H.J., J.L., and B.Y. performed the experiments using the SPID platform. C.C, B.-K.S., J.H.J., and M.B. analyzed all data. C.C. visualized all data. C.C., B.-K.S., M.B., and T.-Y.Y. wrote the manuscript with inputs from all other authors.

## Supplementary Figures

**Fig. S1.**
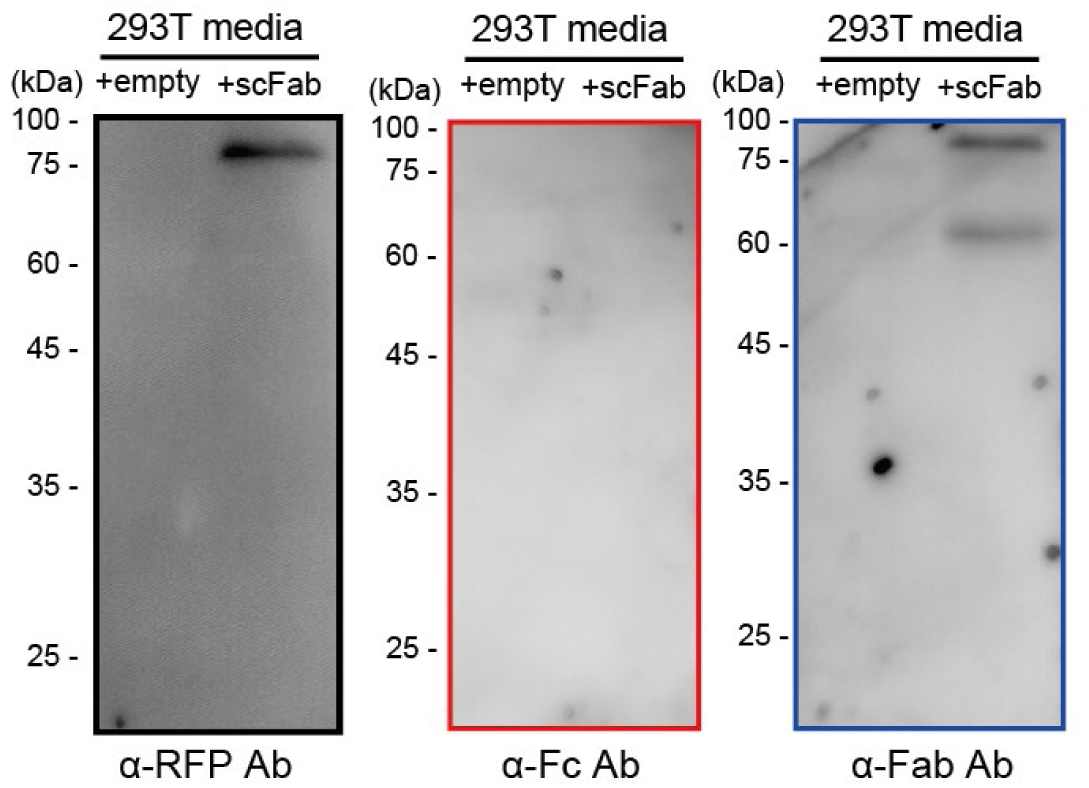
Detection of the secreted Trastuzumab scFabs labeling with RFP in cell culture media by Western blotting.

**Fig. S2.**
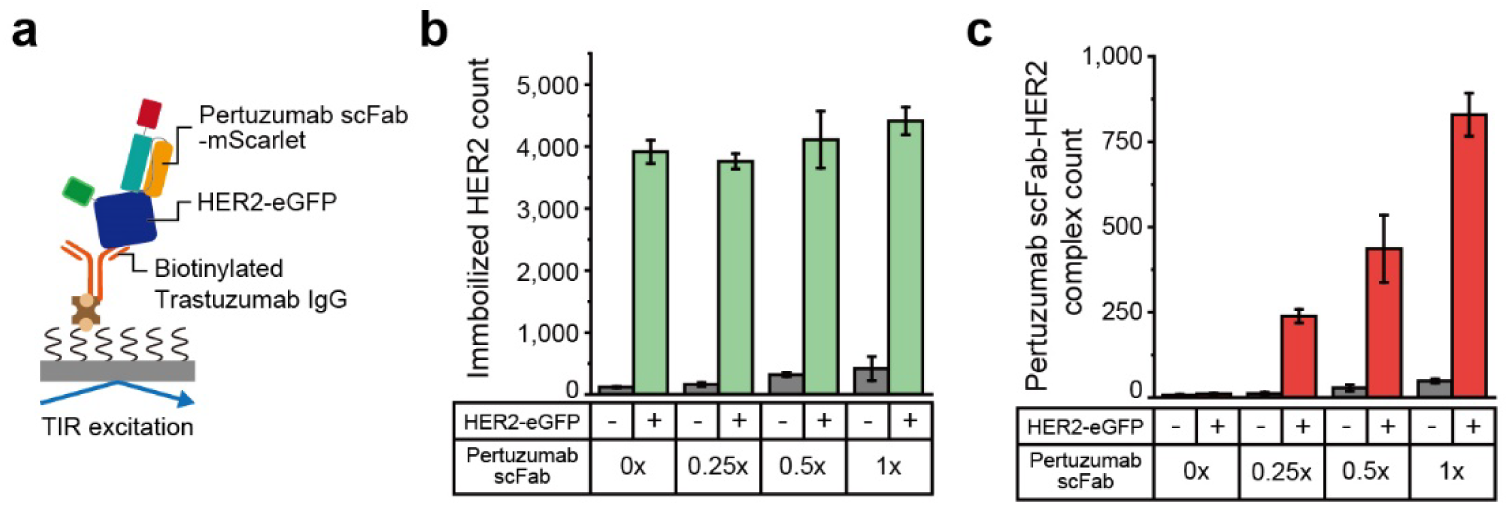
Pertuzumab scFabs can bind to full-length HER2 proteins simultaneously with Trastuzumab. **a**, Schematics for co-immunoprecipitation of Pertuzumab scFab and Trastuzumab IgG. **b,** Surface-immobilization of HER2-eGFP by the biotinylated Trastuzumab IgG using SPID. **c,** Binding of Pertuzumab scFab to HER2-eGFP simultaneously with Trastuzumab IgG.

**Fig. S3.**
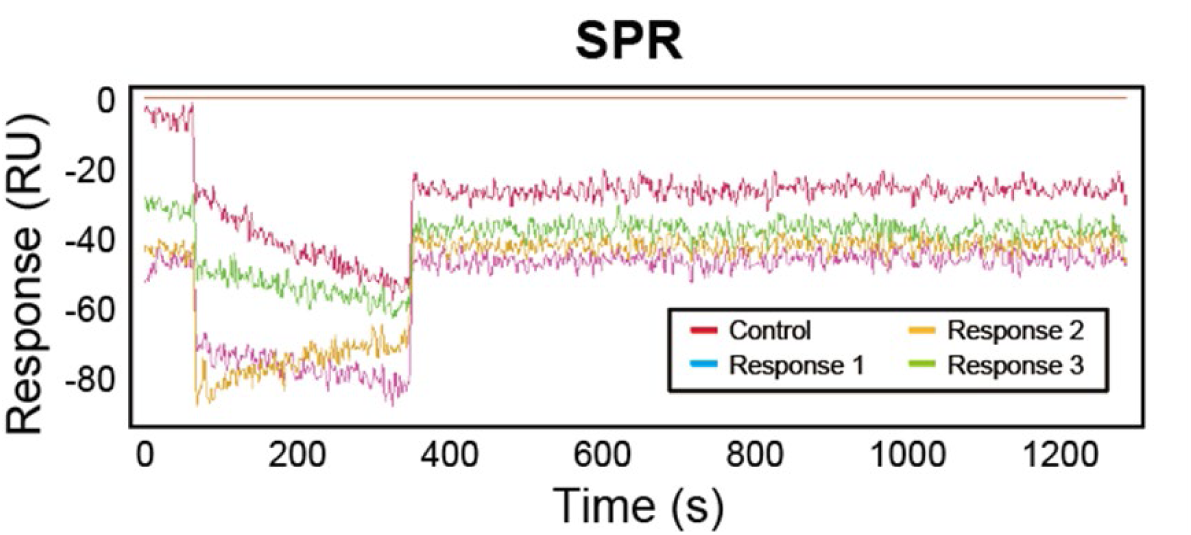
Affinity measurement of Trastuzumab scFabs using HT-SPR. 1,130 pg of crude scFabs and HER2-eGFP antigen lysates failed affinity measurement using HT-SPR.

**Fig. S4.**
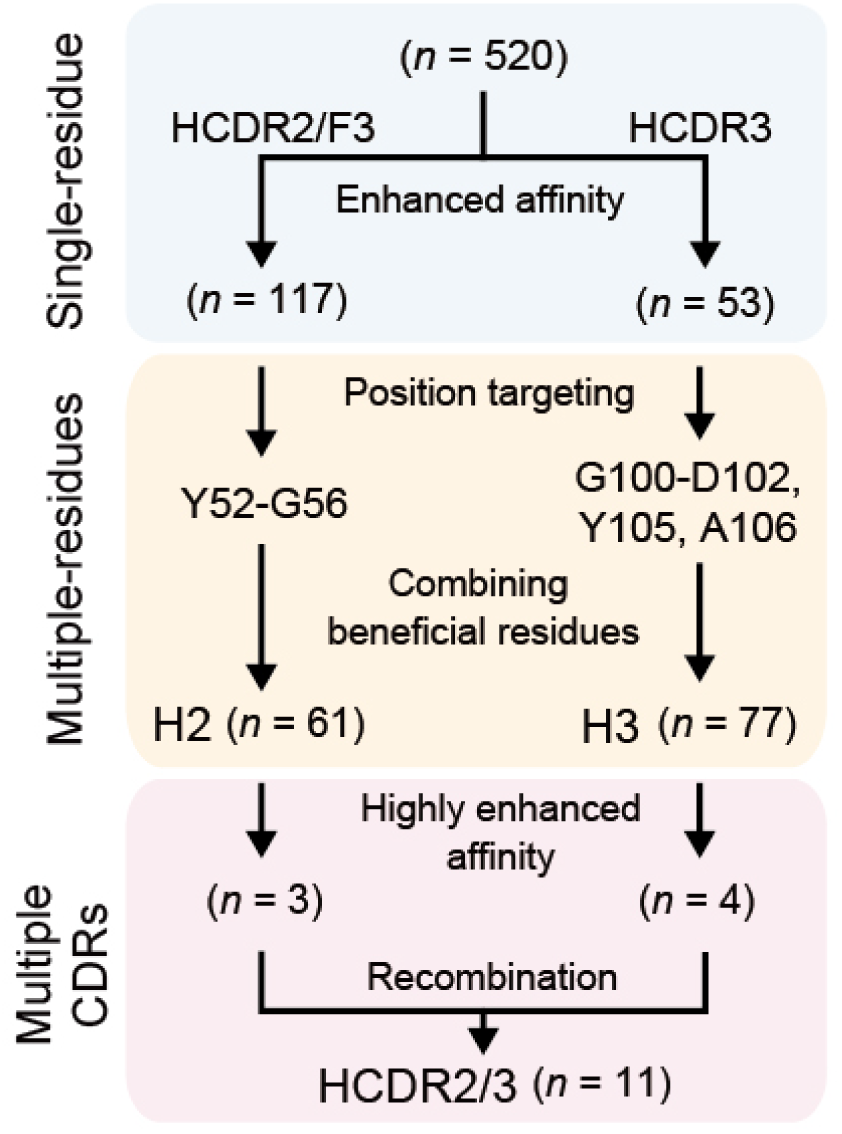
Strategy of systematic engineering toward high-affinity Trastuzumab scFabs.

**Fig. S5.**
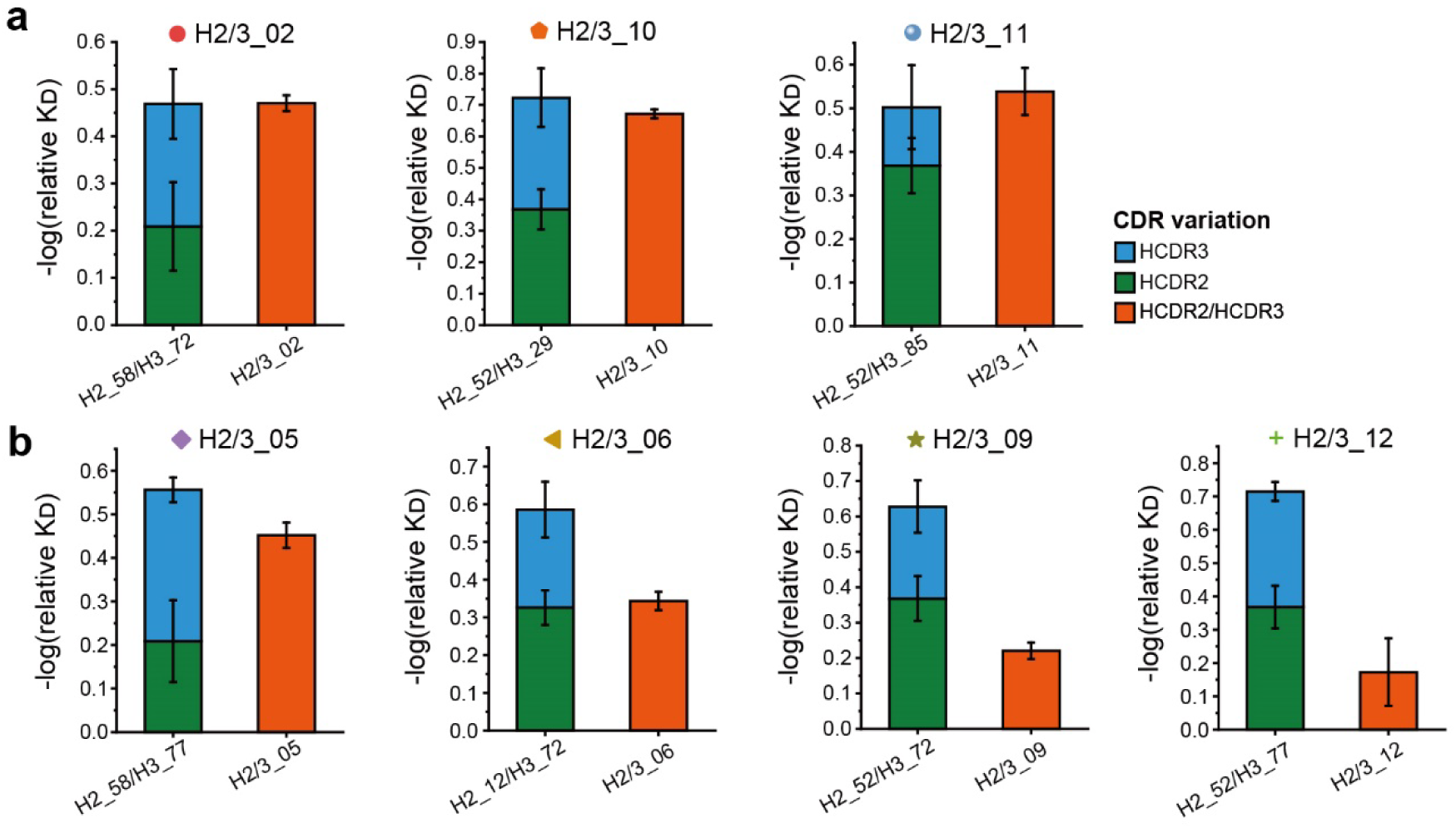
Synergistic analysis for cross-combination of HCDR2 and HCDR3 variants. **a**, Cross-combinatory variants with additive effect. **b,** Cross-combinatory variants with competitive effect.

